# Mammalian N1-adenosine PARylation is a reversible DNA modification

**DOI:** 10.1101/2022.08.17.504244

**Authors:** Michael U. Musheev, Lars Schomacher, Amitava Basu, Dandan Han, Laura Krebs, Carola Scholz, Christof Niehrs

**Affiliations:** Institute of Molecular Biology (IMB), 55128 Mainz, Germany; Division of Molecular Embryology, DKFZ-ZMBH Alliance, 69120 Heidelberg, Germany

## Abstract

Poly-ADP-ribosylation (PARylation) is regarded as a protein-specific modification. However, some PARPs were recently shown to modify DNA termini *in vitro*. Here we use ultrasensitive mass spectrometry (LC-MS/MS), anti-PAR antibodies, and anti-PAR reagents to show that mammalian DNA is physiologically PARylated and to different levels in primary tissues. Inhibition of PAR glycohydrolase (PARG) increases DNA PARylation, supporting that the modification is reversible. DNA PARylation requires PARP1 and *in vitro* PARP1 PARylates single-stranded DNA, while PARG reverts the modification. DNA PARylation occurs at the N1-position of adenosine residues to form N1-Poly(ADP-ribosyl)-deoxyadenosine. Through partial hydrolysis of mammalian gDNA we identify PAR-DNA via the diagnostic deamination product N1-ribosyl-deoxyinosine to occur *in vivo*. We conclude that N1-adenosine PARylation is a novel, reversible DNA modification.

## Main text

ADP-ribosylation of proteins is a posttranslational modification where ADP-ribose from nicotinamide adenine dinucleotide (NAD^+^) is transferred to amino acid residues of target polypeptides. The modification occurs in form of mono- and poly-ADP-ribosylation (MARylation and PARylation, respectively), involving transfer of a single or multiple ADP-ribose moieties, respectively. ADP-ribosylation is catalyzed by the ADP-ribosyltransferase superfamily of enzymes that occur in all kingdoms of life and that are thought to have evolved as a defense mechanism in bacterial virus-host interactions^1^. In mammals, PARylation regulates a vast array of mostly nuclear processes like transcription, DNA replication, chromatin remodelling, RNA splicing, and notably DNA repair^2,3^. PARylation is catalyzed by poly(ADP-ribose) polymerases (PARPs), which generate long branched polymers of ADP-ribose. Among them, PARP1 accounts for most PARylation activity induced by DNA damage, and it is a therapeutic cancer target^**2**,**3**.^

Although in mammals PARylation is regarded as a protein-specific modification, in lower organisms MARylation of DNA was reported. In butterflies, mollusks and *Streptomyces*, the enzyme-toxins pierisin, CARP-1 and Scabin, respectively, can MARylate guanosine residues in DNA^4–6^. In butterflies, DNA-MARylation may potentially serve as defense factor against parasitism^7^. In *Mycobacterium*, the DarT enzyme-toxin MARylates thymidines in single-stranded (ss) DNA^8,9^. Interestingly, it was recently shown *in vitro* that mammalian PARP1 and -2 can PARylate and PARP3 can MARylate phosphorylated DNA 3’- and 5’-ends^10–14^. Thus, DNA PARylation is chemically plausible but whether mammalian DNA is physiologically PARylated, where in the genome the modification occurs, and what its physiological role might be, remains elusive.

### Mammalian DNA is PARylated

We probed PARylation of DNA by dot blot following extensive treatment with Proteinase K to eliminate any residual proteins. We employed a widely used anti-poly ADP-ribose antibody that recognizes linear PAR chains and is free of DNA/RNA cross-reactivity (10H^15^). Total DNA from mouse embryonic stem cell (mESC) and human HEK293T was positive for PAR, while plasmid DNA from *E. coli* (control) was negative (Fig. 1a). We carefully validated the specificity of the antibody signal: i) An anti-pan-ADP-ribose binding reagent (MABE1016) confirmed PARylation of mammalian DNA (Extended Data Figure 1a) and was used hereafter in parallel with 10H anti-PAR antibody; (ii) Treatment with PAR glycohydrolase (PARG), the primary enzyme responsible for degrading cellular ADP-ribose moieties^2^, eliminated the PAR signal of auto-PARylated PARP1 protein (Extended Data Fig. 1b-c) and of mESC DNA (Fig. 1b, Extended Data Fig. 1d), outruling DNA cross-reactivity; iii) DNA PARylation was strongly enhanced when mESCs were grown in presence of PDD, a cell-permeable, inhibitor of PARG^16^ (Fig. 1c, Extended Data Fig. 1e), and reduced in mESCs treated with the PARP inhibitor olaparib17 (Fig. 1d, Extended Data Fig. 1f), supporting anti-PAR specificity; iv) DNase I digestion erased the PAR signal, indicating that it did not stem from residual proteins or co-purifying free PAR chains released by Proteinase K digestion (Extended Data Fig. 1g). On the other hand, DNase I treatment did not affect the PAR signal of free PAR chains, ruling out that DNase I may degrade PAR chains directly (Extended Data Fig. 1g); v) Combined RNase A, RNase H, and RNase III treatment had no effect on the PAR signal from genomic DNA (Extended Data Fig. 1h), ruling out PARylation of contaminating RNA18; vi) Immunoprecipitation of mESC DNA using an anti-dsDNA antibody enriched for PAR chains (Fig. 1e); vii) Southwestern blot analysis of mESC DNA with anti-PAR antibody (Fig. 1f) and HEK293T DNA with both anti-PAR antibody and anti-pan-ADPr reagent (Extended Data Figure 1i-j) yielded a signal in form of a smear only after restriction digest, indicating that PAR chains are covalently attached to high molecular weight DNA, instead of free or protein-bound PAR chains co-purifying during DNA preparation. The average molecular weight of the PAR signal smear was expectedly higher for the 6-base cutter EcoRI than for the 4-base cutter MseI. Furthermore, the PAR signal was strongly reduced by treatment with PARG but not by a second Proteinase K digest/phenol-chloroform purification (Fig. 1f); viii) Dot blot analysis of DNA from diverse human organs revealed PARylation in all tested tissues but with notable differences (Extended Data Fig. 1k), suggesting biological specificity.

**Fig. 1:**
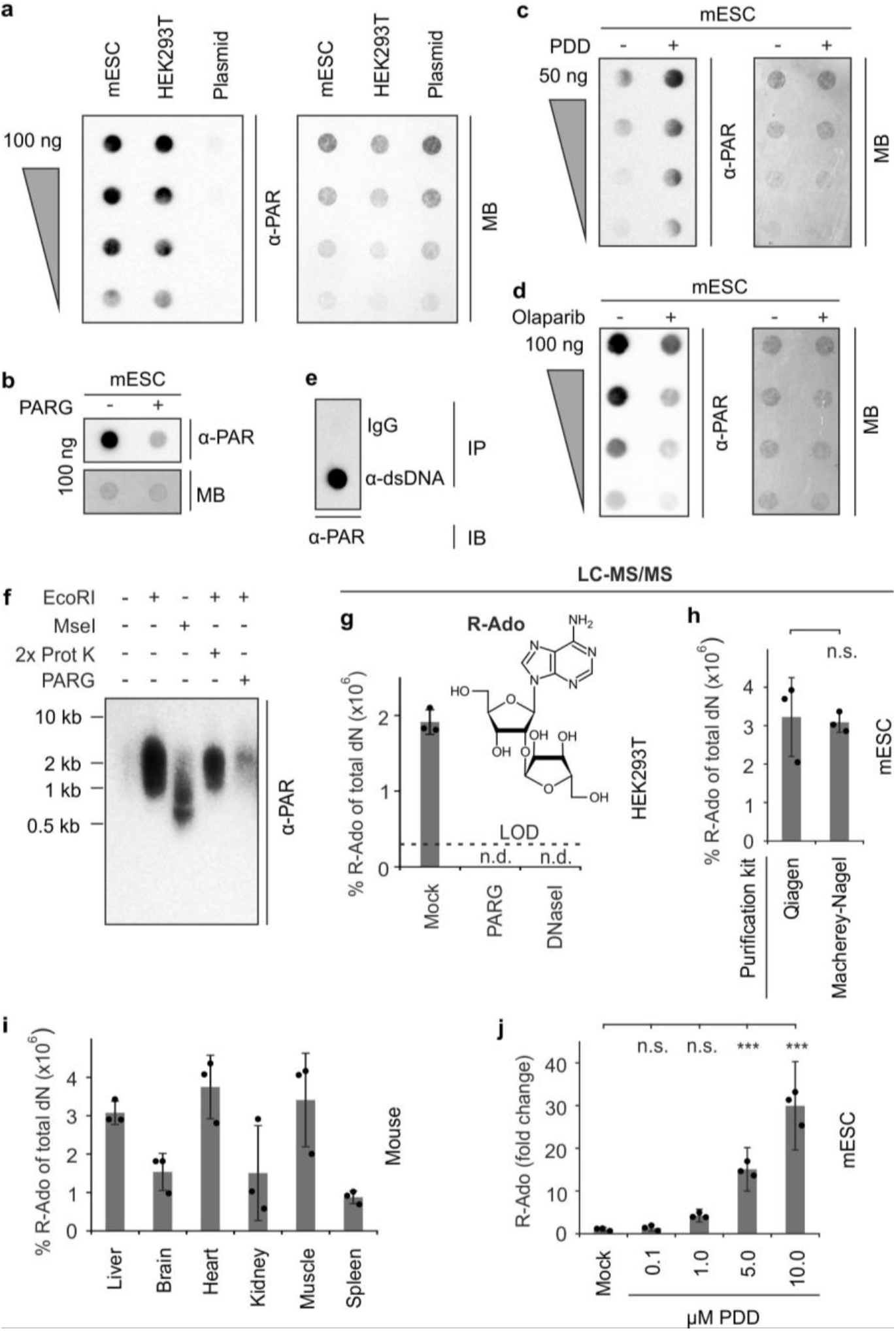
Mouse and human genomic DNA is PARylated. **a-e**, Dot blot analysis for PARylation of mESC, HEK293T and plasmid DNA (a), mESC DNA treated with PARG (**b**), DNA purified from mESCs treated with the PARG inhibitor PDD00017273 (**c**), DNA purified from mESCs treated with the PARP inhibitor olaparib (**d**), genomic mESC DNA after IP with IgG or anti-dsDNA antibody (**e**). gDNA was serially diluted (2x) in a and c-d, MB, methylene blue. **f**, Southwestern blot analysis for PARylation of mESC DNA treated with EcoRI, MseI, Proteinase K, and PARG as indicated. Length of marker DNA is shown on the left. **g**, LC-MS/MS quantification of ribosyl-adenosine (R-Ado) on HEK293T DNA treated as indicated. Samples were repurified after enzyme treatments by a second column-based DNA purification to remove any PAR and DNA monomers (dashed line, limit of detection (LOD); s.d., n = 3 biological replicates, n.d., not detected). **h-j**, LC-MS/MS quantification of R-Ado of mESC DNA isolated by two different DNA preparation kits (Qiagen Blood & Cell culture DNA kit or Macherey-Nagel NucleoBond HMW DNA kit; s.d., n = 3 biological replicates; n.s., not significant by two-sided, unpaired t-test for unequal variances, **h**), of DNA from the indicated adult female mouse tissues (s.d., n = biological replicates from 3 individual mice, **i**), of DNA from mESCs treated with increasing amounts of the PARG inhibitor PDD00017273. R-Ado levels from mock-treatment is arbitrarily set to 1. s.d., n = 3 biological replicates; \*\*\**P* < 0.005, not significant (n.s.) by Dunnett’s test, **j**.

To confirm mammalian DNA-PARylation antibody-independently, we employed stable isotope dilution mass spectrometry (LC-MS/MS), the gold standard in quantification of modified nucleosides. DNA from HEK293T was enzymatically digested to nucleosides and analyzed by LC-MS/MS for ribosyl-adenosine (R-Ado), a product diagnostic for linear PAR chains^19^. R-Ado analysis was highly sensitive and accurate, with a limit of detection (LOD) of 3× 10^−17^ moles, or ∼6 R-Ado molecules per mammalian genome. In DNA from HEK293T cells, R-Ado was detectable at very low levels of 2× 10^−6^ % of total dN, corresponding to ∼100 R-Ado monomers per genome (Fig. 1g). Again, PARG or DNaseI treatment reduced R-Ado signals below detection limit. R-Ado was also detected in DNA from mESCs and mouse organs up to 4× 10^−6^ % of total dN (Fig. 1h-i). Given that PAR chains can have a chain length in the order of ∼100 residues^20^, this yields an average of 1-2 PARylation sites per genome. Notably, treating mESCs with increasing doses of the PARG inhibitor PDD increased DNA PARylation up to 30-fold, indicating reversibility and high turnover of the modification (Fig. 1j).

We conclude that mammalian PARylation is a rare and reversible DNA modification occurring in primary mammalian tissues and cell lines.

### DNA PARylation requires PARP1 and is unrelated to DNA breaks

Which enzyme might PARylate DNA in mammals? PARylation is catalyzed by the PARP family that consists of 17 and 16 members in human and mouse, respectively^21^. Based on their expression level in mESCs and HEK293T cells, we selected PARP1, PARP2, PARP4, TNKS (PARP5A) TNKS2 (PARP5B), PARP7 and ZC3HAV1 (PARP13) as candidates for further analysis (Extended Data Fig. 2a-b). In HEK293T cells, we observed a tendency for siRNA-mediated knockdown of *PARP1* but not the other tested *PARPs* to decrease global DNA PARylation while in mESCs both *Parp1* and *Parp2* knockdowns significantly decreased DNA PARylation measured by LC-MS/MS quantification of R-Ado (Fig. 2a-b).

**Fig. 2:**
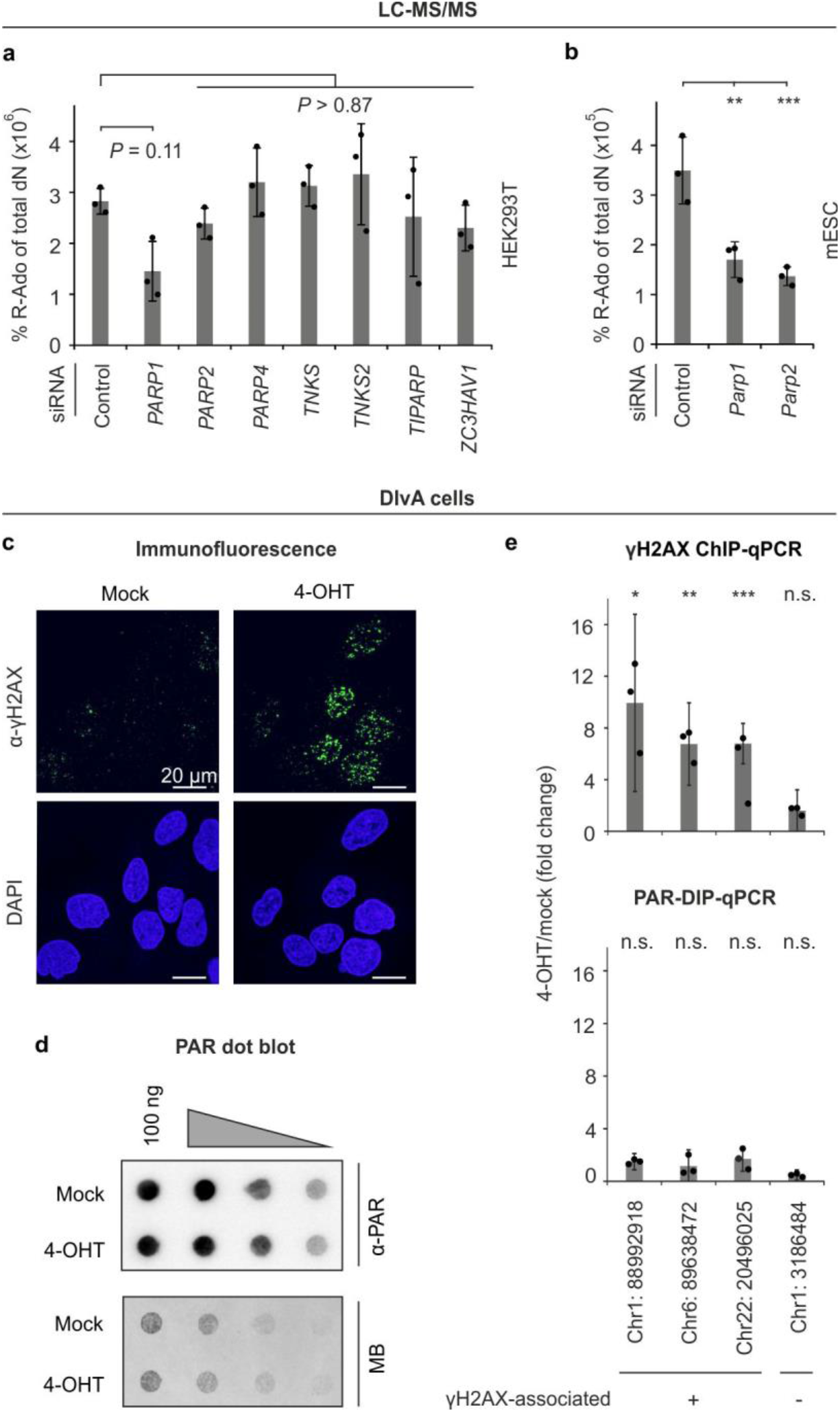
DNA PARylation requires PARP1 but is not triggered by strand breaks *in vivo*. **a-b**, LC-MS/MS quantification of ribosyladenosine (R-Ado) in 293T (**a**) and mESC (**b**) DNA after siRNA depletion of selected *PARPs* (s.d., n = 3 biological replicates). \*\**P* < 0.01, \*\*\**P* < 0.005, and indicated p-values as by Dunnett’s test. **c**, γH2AX immunofluorescence of DIvA cells in which AsiSI nuclease expression was induced by 4-OHT treatment. DNA is stained with DAPI. **d**, Dot blot analysis for PARylation of DNA from AsiSI nuclease expressing DIvA cells treated as in (f) with serially diluted (2x) DNA. **e**, Left: γH2AX ChIP-qPCR analysis of selected genomic loci DIvA cells previously described to be associated with (+) or without (-) γH2AX upon 4-OHT treatment to induce AsiSI nuclease24. Right: PAR-DIP-qPCR analysis of the same loci as shown on the left. Target regions are named by chromosomal location of the AsiSI recognition site in the human hg38 reference assembly. s.d., n = 3 biological replicates; \**P* < 0.05, \*\**P* < 0.01, \*\*\**P* < 0.005, not significant (n.s.) over mock by two-sided, unpaired t-test for unequal variances.

PARP1 plays a key role in the DNA damage response, where it recognizes DNA strand breaks and PARylates nearby proteins to initiate repair^22^. In addition, PARP proteins can PARylate DNA ends *in vitro* and it was proposed that this may reflect a physiological role during break repair^23^. Hence, we examined whether DNA PARylation may generally localize to DNA strand breaks (DSB). We made use of DIvA cells as experimental cell system, which harbor the DNA rare cutter AsiSI that in presence of 4-hydroxytamoxifen (4-OHT) creates DSBs at defined positions across the genome and in various chromatin contexts^24^. While 4-OHT treatment of DIvA cells robustly induced gH2AX nuclear foci (Fig. 2c), there was no increase in global DNA PARylation (Fig. 2d). To specifically monitor local changes of DNA PARylation, we performed PAR-DIP-qPCR for three known AsiSI-mediated DNA break loci^24^. While 4-OHT-induced γH2AX recruitment to these loci, PAR signals were low and remained unchanged by 4-OHT (Fig. 2e). We conclude that DNA PARylation is not a general feature of DNA breaks.

### DNA PARylation by PARP1 occurs at adenosine residues in single-stranded DNA

DNA-PARylation, if uncontrolled, may be deleterious to cells, so there must be mechanisms that provide specificity. PARP1 may display substrate specificity towards certain types of DNA. To analyze substrate specificity, we performed *in vitro* PARylation reactions with PARP1 (Extended Data Fig. 3a) and an arbitrary 30 nt DNA-oligonucleotide (hereafter referred as ‘standard oligo’) carrying a ^32^P-end-labeled 5’-phosphate. We tested the standard oligo as ssDNA and dsDNA with either blunt-or 3’-recessed end (Fig. 3a). Reaction products were analyzed by denaturing gel electrophoresis, where PARylated DNA accumulates at the top of the gel^13^. Interestingly, ssDNA was robustly PARylated, unlike hybrid DNA (Fig. 3b). The reaction required the PARP1 co-substrate NAD^+^ and was inhibited by the PARP-inhibitor olaparib (Fig. 3c). DNA PARyation was not due to spontaneous-or PARP1-catalyzed addition of pre-existing PAR chains to the standard oligo (Extended Data Fig. 3b) and PARP1 did not PARylate ssRNA (Extended Data Fig. 3c-d)

**Fig. 3:**
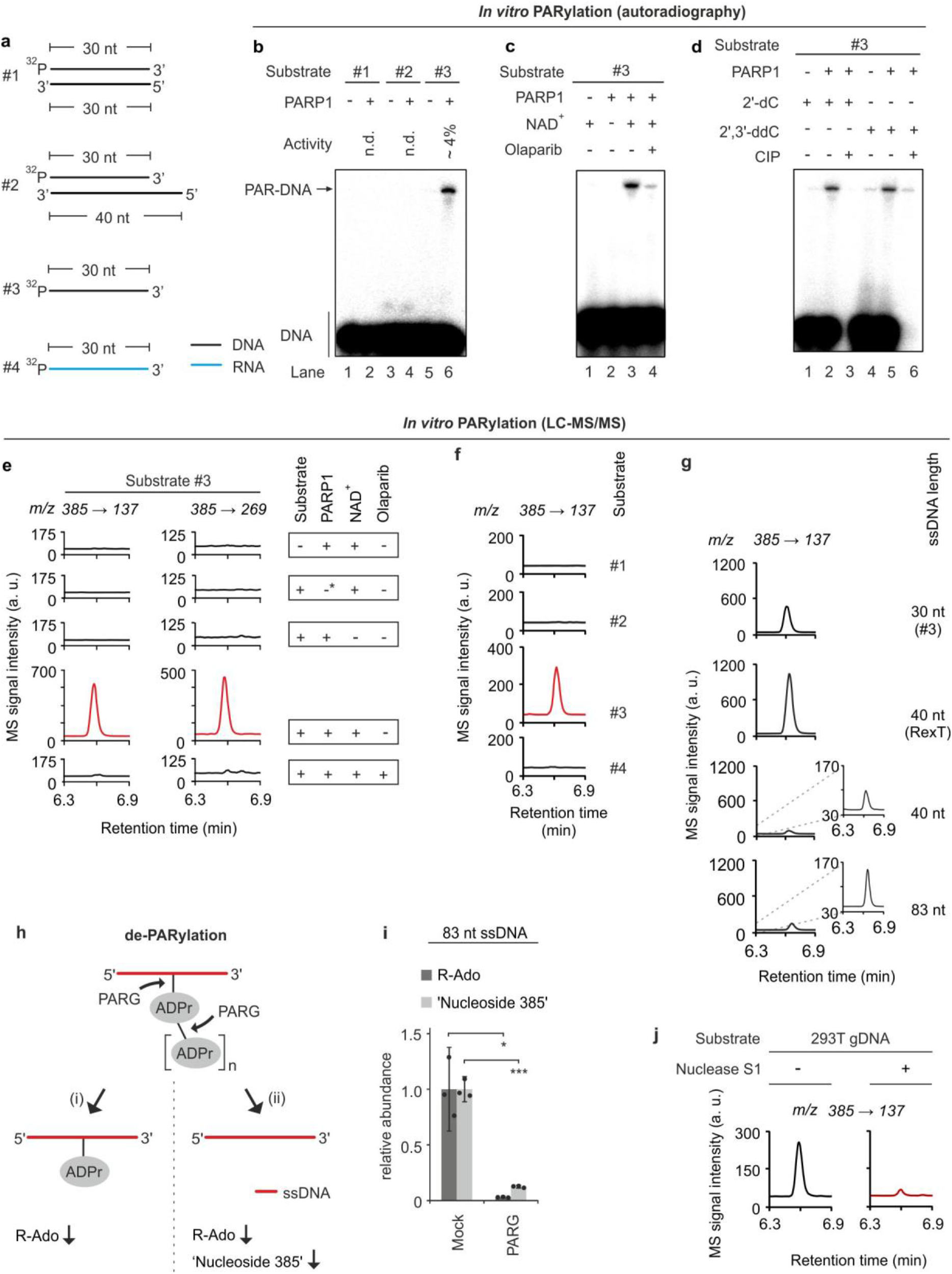
PARP1 PARylates ssDNA at a nucleobase. **a**, Scheme of substrates #1 - #4 used for *in vitro* PARylation assays. The 30 nt black strand is designated as standard oligo in the main text. 32P, 5’-phosphate with 32P-radiolabel. **b-d**, Autoradiography of denaturing PAGE of reaction products from *in vitro* PARylation assay with substrates shown in (**a**) in presence or absence of human PARP1, NAD+ and the PARP1 inhibitor olaparib. In (**d**), substrate #3 contained either a 2’-dC or a 2’,3’-ddC terminal nucleotide and was treated with or without calf intestine phosphatase (CIP) post reaction. **e-g**, LC-MS/MS chromatograms of reaction products from *in vitro* PARylation assays in presence of native or heat-denatured PARP1: (**e**), with or without substrate #3, NAD+ and olaparib as indicated. Left and right panels show mass transitions corresponding to mass shifts expected for loss of a deoxyribose + ribose (*m/z* 385 → 137) and loss of a single deoxyribose (*m/z* 385 → 269) from the parental ‘nucleoside 385’. -*, heat-denatured PARP1. (**f**), with substrates #1 - #4 as shown in (a). Mass transition is shown as expected for loss of a deoxyribose + ribose (*m/z* 385 → 137) from the parental molecule ‘nucleoside 385’. (**g**), with standard substrate #3, and three additional ssDNA oligonucleotides (RexT13, 40 nt, 83 nt). The signal of the lower two chromatograms is magnified on the right.**h**, Scheme for two alternative outcomes of de-PARylation of PARylated DNA with PARG. (i) PARG hydrolyses the ADP ribose polymer but not the terminal ADP ribose unit at the DNA base acceptor side. LC-MS/MS analysis of the reaction products would yield decreased R-Ado and unchanged ‘nucleoside 385’ levels. (ii) In addition to polymer degradation, PARG cleaves the linkage of the ADP ribose and the DNA base acceptor side resulting in both decreased R-Ado and ‘nucleoside 385’ levels. The latter outcome would indicate complete reversibility of DNA base PARylation. **i**, LC-MS/MS quantification of R-Ado and ‘nucleoside 385’ on the 83mer ssDNA oligo (see panel g) after *in vitro* PARylation by PARP1 followed by mock or PARG treatment of the reaction products as indicated. s.d., n = 3 replicates; \**P* < 0.05 and \*\*\**P* < 0.005 by two-sided, unpaired t-test for unequal variances. Note reduced ‘nucleoside 385’ levels after PARG treatment, which supports complete reversibility of DNA base PARylation by PARG. **j**, PARP1 forms ‘nucleoside 385’ with single stranded genomic DNA. LC-MS/MS chromatograms for ‘nucleoside 385’ of reaction products from *in vitro* PARylation. Genomic DNA from 293T cells pretreated or not with Nuclease S1 was incubated with PARP1 and NAD+.

Since PARP1 was reported to PARylate both 5′- and 3′-terminal phosphates of complex DSB-mimicking oligonucleotides *in vitro* ^10,13^, we released terminal ^32^P-end-labeled 5’-phosphate with calf intestine phosphatase (CIP). If PARylation occurs on the labeled 5’-phosphate, the modification should protect the label against CIP treatment. However, CIP treatment erased the signal of both the unmodified and PARylated standard oligo, thus excluding PARylation of the DNA 5’-phosphate terminus (Fig. 3d). Conversely, the 3’-end was also unmodified, since PARylation occurred with an oligo terminating in a dideoxy-cytosine (2’3’-ddC), i.e. lacking a 3’-OH acceptor site. As control, the PAR signal still decreased upon CIP treatment, ruling out alternating PAR-acceptor sites (Fig. 3d). These results suggest that in ssDNA, PARylation occurs at DNA bases rather than at termini. We also tested an oligo (RexT) previously shown to be 5’-terminal PARylated by PARP1^13^. Following *in vitro* PARylation, CIP treatment removed ∼ 70% of the signal, confirming that while there is 5’-phosphate modification, PARylation mostly occurs internally (Extended Data Fig. 3e).

To identify the PAR acceptor site on ssDNA, we adopted a method^19^ that enzymatically degrades the PAR chains and leaves a single ribose remnant on the PAR acceptor molecule. Conveniently, this procedure also degrades DNA to single nucleosides. Consequently, we degraded the *in vitro* PARylated standard oligo and screened the products by LC-MS/MS for ribosylated nucleosides, i.e. containing both a ribose moiety remaining from PARylation and a deoxyribose. Intriguingly, we identified a positively charged ion with a mass over charge ratio (*m/z*) of 385 (‘nucleoside 385’) and mass transitions that fit to loss of a ribose and a deoxyribose (*m/z* = 385 → 137), as well as loss of a single deoxyribose (*m/z* = 385 → 269), supporting the presence of both sugars on the nucleobase (Fig. 3e). Generation of ‘nucleoside 385’ was dependent on active PARP1, the standard oligo, NAD^+^, and was sensitive to olaparib treatment. We observed no signals for ‘nucleoside 385’ from dsDNA or ssRNA after PARP1 incubation (Fig. 3f). Thus, levels of ‘nucleoside 385’ closely correlate with levels of radiolabeled PARylation products (Fig. 3b-c, Extended Data Fig. 3c). ‘Nucleoside 385’ was also formed with varying efficiency using three additional ssDNA oligos of different sequences and lengths, including RexT13 (Fig. 3g). PARG treatment of a PARylated 83mer ssDNA strongly reduced the levels of ‘nucleoside 385’, indicative of reversibility of base PARylation *in vitro* (Fig. 3h-i). Furthermore, we detected ‘nucleoside 385’ when using HEK293T genomic DNA instead of synthetic DNA oligonucleotides as substrate for PARP1 *in vitro* PARylation. Again, the reaction was ssDNA-dependent as pre-incubation of gDNA with Nuclease S1 prevented ‘nucleoside 385’ formation (Fig. 3j).

To identify the ribosylated base, we first used ssDNA oligos homopolymeric for all four canonical nucleosides but could not detect ‘nucleoside 385’ after *in vitro* PARylation (Extended Data Fig. 4a). The result suggests a requirement of base PARylation for certain sequence contexts and/or secondary structures in ssDNA. Next, we *in vitro*-PARylated four 83mer ssDNA substrates with PARP1, each containing one isotopically labelled nucleotide (^15^N_5_-dG, ^15^N_3_-dC, ^15^N_2_-T or ^13^C_10_-dA, respectively), and scanned the reaction products for the mass of a respective heavy ‘nucleoside 385’. The only reaction that yielded the expected mass shift of ‘nucleoside 385’ was the ^13^C_10_-dA 83mer oligo, indicating adenine as the PAR acceptor base (Fig. 4a). Interestingly, replacing carbon-labeled ^13^C_10_-dA by nitrogen-labeled ^15^N_5_-dA on the 83mer substrate, we could not detect the expected ‘nucleoside 385’ + 5 Da = 390 ion product but instead a ‘nucleoside 385’ + 4 Da = 389 ion product (Fig. 4b). The result indicates that the ‘nucleoside 385’ adenosine derivative lost one labeled nitrogen atom before, during, or after PARylation. Indeed, adenosine can lose a nitrogen by deamination to inosine25. Strikingly, the predicted *m/z* for ionized N1-ribosyl-deoxyinosine (M_r_ + H^+^) matches exactly the observed *m/z* of ‘nucleoside 385’. Hence, we first reasoned that inosine formed by deamination of adenosine might be the true acceptor residue for PARylation. We therefore *in vitro* PARylated the standard 30 nt oligo, in which all six dA residues were substituted by dI (Extended Data Fig. 4b). However, we detected neither significant ‘nucleoside 385’ by LC-MS/MS nor any PARylated oligo in radioactive PARylation assay (Extended Data Fig. 4c-d) excluding inosine as acceptor for PARylation. We next considered that deamination might not be cause but consequence of PARylation of dA, i.e. ribosylated dA might be prone to deamination. While deamination of adenosine is slow in intact DNA26, N1-substituted dA adducts readily deaminate27,28. Hence, we hypothesized that N1-ribosyl-dA is the precursor of N1-ribosyl-dI (Extended Data Fig. 4e). However, the deaminase inhibitor pentostatin added to the PARylation reaction as well as to the enzymatic degradation step had no effect on the levels of ‘nucleoside 385’ or the occurrence of a signal expected for N1-ribosyl-dA (Extended Data Fig. 4f). This result argues against enzymatic deamination by contaminating deaminases and points towards spontaneous deamination. To reduce spontaneous deamination, we omitted a 95 °C denaturation step in the DNA processing protocol and avoided prolonged sample storage prior to LC-MS/MS analysis. Using these precautions, we indeed detected in the PAR reaction of the standard oligo besides ‘nucleoside 385’ a new signal with an *m/z* matching N1-ribosyl-dA (*m/z* = 384 → 136) (Fig. 4c-d). Moreover, when we analyzed under these conditions a ^15^N_5_-dA-labeled 83mer DNA PARylation product, we detected two signals with the same *m/z* transition but different LC-retention times, as expected for 15N_5_-labeled N1-ribosyl-dA and ^15^N_4_-labeled N1-ribosyl-dI (Extended Data Fig. 4g-h). Further analysis revealed that deamination is slow on PARylated-dA in intact DNA but accelerated on nucleosides (Extended Data Fig. 5a-c). We conclude that PARP1 catalyzes PARylation of ssDNA on adenosine-N1 forming N1-Poly(ADP-ribosyl)-dA.

**Fig. 4:**
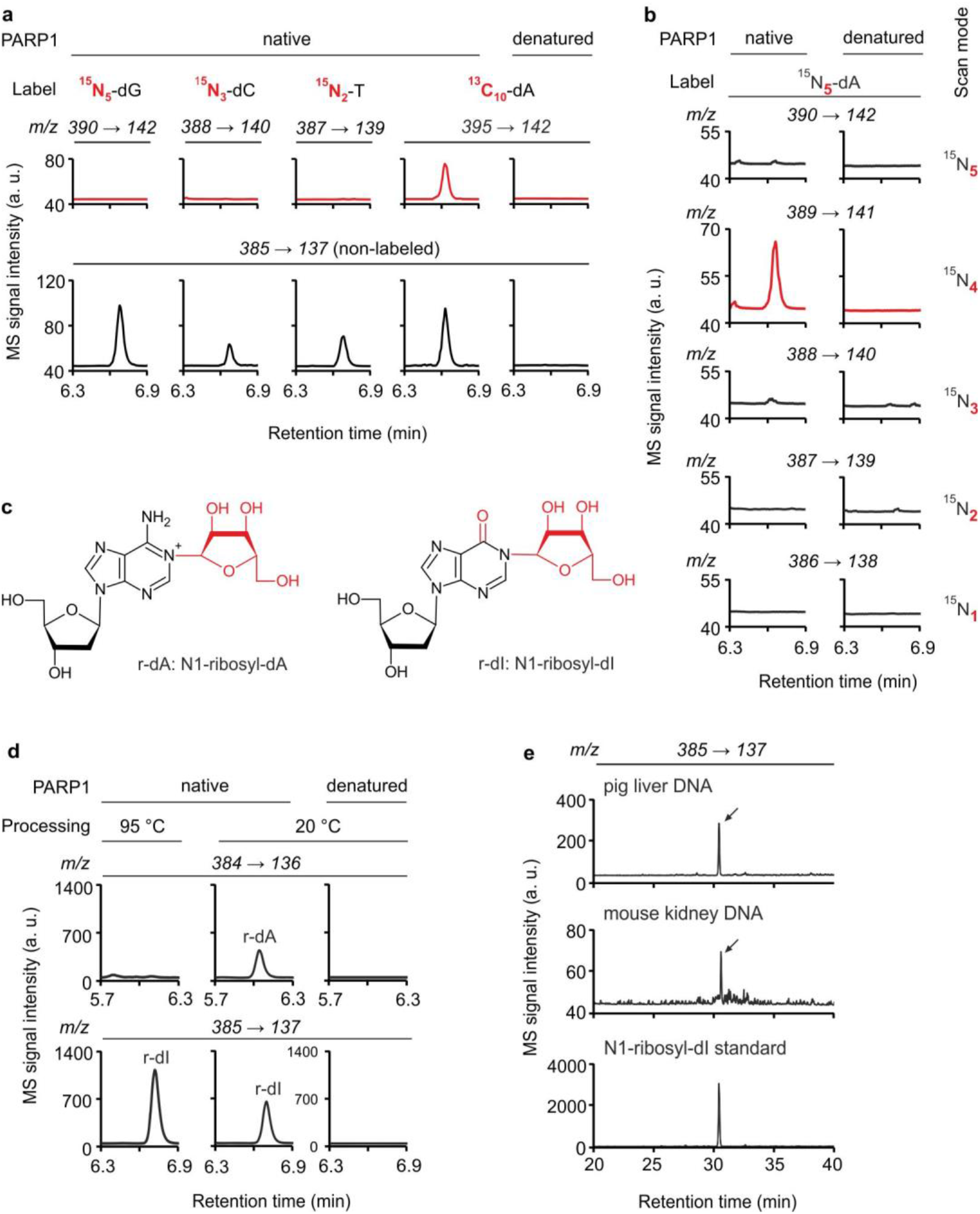
DNA PARylation occurs at the N1-position of adenosine residues *in vitro* and *in vivo*. **a**, LC-MS/MS chromatograms of reaction products from *in vitro* PARylation assays in presence of native or heat-denatured PARP1 with four 83mer ssDNA substrates each bearing one type of heavy isotope-labeled nucleoside as indicated. Top, scan for signals with *m/z* transitions expected for a mass shift of ‘nucleoside 385’ from each label. Bottom, detection of signals that correspond to non-labeled ‘nucleoside 385’. Note, all four substrates contain a mixture of the respective labeled and unlabeled nucleoside. **b**, LC-MS/MS chromatograms of reaction products from *in vitro* PARylation assays as in (a) but with a 15N_5_-dA-labeled 83mer ssDNA substrate. Reaction products were scanned for signals with *m/z* transitions that correspond to mass shifts of +5 to +1 of ‘nucleoside 385’ (from top to bottom). **c**, Spontaneous deamination of N1-ribosyl dA (r-dA, attached ribose moiety in red) leads to N1-ribosyl-dI (r-dI, ribose moiety and O6 in red). **d**, LC-MS/MS chromatograms of reaction products from *in vitro* PARylation assays as in (a) but with substrate #3 in which reaction products were either processed at 95 °C or 20 °C before mass spec analysis, and screened for signals with *m/z* transitions expected for N1-ribosyl-dA (r-dA, *m/z* 384 → 136, top) or N1-ribosyl-dI (r-dI, *m/z* 385 → 137, bottom). **e**, LC-MS/MS chromatograms of N1-ribosyl-dI, (arrows, *m/z* 385 → 137) enriched from 5 mg pig liver and 2.5 mg mouse kidney DNA and compared to the reaction product of an *in vitro* PARylation assay with PARP1 on substrate #3.

To demonstrate that DNA PARylation occurs on N1-adenosine *in vivo*, we developed an LC-MS/MS protocol to detect in hydrolyzed gDNA the diagnostic deamination product, N1-ribosyl-dI. Expecting it to be ultra-rare, we used several milligrams of pig liver and mouse kidney DNA.

Separating ssDNA from dsDNA and enriching by HPLC, we indeed detected N1-ribosyl-dI in DNA from both tissues (Fig. 4e).

## Discussion

PARylation is an important posttranslational modification of proteins that is involved in numerous processes. In mammalian cells, PARylation is thought to be limited to proteins. PARylation of mammalian DNA therefore comes as a surprise. While rare in abundance, DNA PARylation is widespread in mammalian DNA from different cell lines, tissues, and species. We provide evidence that DNA PARylation by PARP1 occurs on adenosine N1, which is indeed the most nucleophilic atom of adenine^29^, providing a rationale for its PARylation. We also show that PARP1 specifically acts on ssDNA, likely due to greater accessibility of unpaired adenosine residues, because the adenine N1 position is engaged in dsDNA base-pairing. Certain bacterial, insect and mollusc mono-ADP-ribosyltransferases are also known to modify ssDNA and dsDNA on thymidine and on the N2 position of guanosine^4–6,8,9,^ but modification of adenosine has, to our knowledge, not been described.

We show that PARylation requires PARP1 and PARP2, two major PARP enzymes that have been widely studied in protein PARylation^3^. Indeed, PARP1 is known to recognize distortions in the DNA helical backbone and its enzymatic activity is activated by hairpins, cruciforms, and stably unpaired regions in double-stranded DNA^30^. Moreover, PARP1 is involved in various processes that also involve single-stranded DNA, e.g. replication, transcription, or DNA damage repair^22^. However, we show that DNA PARylation is likely not a generalized feature of DNA breaks. Results from *in vitro* DNA PARylation by PARP1 support that sequence contexts or secondary structure of ssDNA are essential. Furthermore, given the low efficiency of DNA PARylation we observe *in vitro*, PARP1 *in vivo* may also employ cofactors^31,32^ that enhance the reaction and provide additional specificity.

PARylation of proteins is reversible by PAR-glycohydrolases^33^ and we provide evidence that DNA PARylation is also reversible *in vitro* and *in vivo* by PARG, indicating that it is dynamic and subject to regulation. However, since the PARG inhibitor PDD that we employed is not selective for PARG^34^, other glycohydrolases may also reverse DNA PARylation *in vivo*, including TARG1, MACROD1/2 and ARH3^33^.

A key open question that remains to be addressed is where in the genome PARylation occurs. The prediction of our *in vitro* PARylation assays is that the PARP1 target is single stranded DNA, which physiologically occurs in different contexts, for example in DNA:RNA hybrids (R-loops), replicating DNA, DNA repair intermediates, G-quadruplex DNA, cDNA, and DNA tandem repeats. To map DNA PARylation genome-wide, we attempted PARylated DNA immunoprecipitation followed by next generation sequencing (PAR-DIP-seq). However, standard sample workup during next generation sequencing precludes amplification and detection of the PARylated DNA strand because of steric hindrance by PAR chains that stall DNA polymerases. Thus, methods need to be developed that overcome this technical challenge in the future. PAR-DIP-seq will then allow addressing the physiological substrate specificity and the biological relevance of this novel mammalian DNA modification.

Another challenge in addressing the biological function of PARylation by PARP1 is its pleiotropy, the vast number of substrates and processes in which this protein is involved, including transcription and DNA repair^22^, which render it difficult to interpret loss-of-function effects and establish causality. Hence, a separation-of-function PARP1 mutant that is deficient in PARylation of DNA but not of protein will be an ideal tool to address the physiological role of DNA *vs*. protein PARylation in the future.

Our study establishes the conceptual and methodological framework to address these and other emerging questions.

## Acknowledgements

We thank Jean-Christophe Amé, Jef Boeke, Meelad M. Dawlaty, Gaëlle Legube, and Robert Rümmler for materials, and Joan Barau for critical reading of the manuscript. We gratefully acknowledge technical support and advice by the IMB core facilities Protein Production, Bioinformatics, and Microscopy and their members Martin Möckel, Fridolin Kielisch, and Sandra Ritz. This work was supported by the Deutsche Forschungsgemeinschaft (DFG, German Research Foundation) instrument funding – [INST247/768-1 FUGG] and the ERC Advanced Grant *HybReader*.

## Author Contributions

Conceptualization, M.U.M., L.S., A.B. and C.N.; Methodology, M.U.M., L.S., A.B., D.H., L.K., C.S.; Investigation, M.U.M., L.S. and A.B.; Writing – Original Draft, L.S. and C.N..; Review and Editing, M.U.M., L.S., A.B. and C.N.; Supervision, L.S. and C.N.; Funding Acquisition C.N.

## Corresponding authors

Correspondence to Christof Niehrs or Lars Schomacher.

