## Supplementary data for "Mammalian N1-adenosine PARylation is a reversible DNA modification"

### Methods

#### Cell culture and transfection

Mouse embryonic stem cell (mESC) clone WT #4[35] was cultured on tissue culture plates coated with 0.1% Gelatin (Millipore) in 2i medium (Neurobasal - DMEM/F-12 medium (Gibco), supplemented with 1x N2 (Gibco), 1x B27 (Gibco), 2 mM L-Glutamine (Gibco), 1000 U/ml Leukemia inhibitory factor (LIF, Millipore), 100 U/ml PEN-STREP (Gibco), 1 µM PD0325901 (Sigma), 3 µM CHIR99021 (Sigma), 50 µg/ml BSA) at 37°C in 5% $CO_2$ and 20% $O_2$. HEK293T (ATCC number CRL-11268) and DIvA cells[24] (kind gift from G. Legube) were cultured in DMEM (Gibco), K562 cells (ATCC number CCL243) in RPMI (Gibco) both supplemented with 10% FBS Gold (PAA), 2 mM L-glutamine, and 100 U/ml PEN-STREP at 37 °C in 5% CO2 and 20% $O_2$. All cell lines were tested negative for mycoplasma contamination.

Cells were transiently transfected with siRNAs and plasmid DNA with Lipofectamine 2000 (Life Technologies) according to the manufacturer’s instructions. siRNA knockdown efficiencies were routinely assessed by reverse transcription (RT)-coupled qPCR, and were at least 60% efficient. For PARG inhibition, mESCs were treated with 0.1 - 10 µM PDD00017273 (Sigma) for 48 h. PARP inhibititon was for 24 h with 1 µM olaparib (Selleckchem, S1060).

#### Protein purification

*PARG*

Plasmid pGEX6P3-hPARGfl[36] encoding a codon-optimized full-length human poly (ADP-ribose) glycohydrolase (PARG) was a gift from J.C. Amé. Expression and purification of PARG was essentially as described before[36] using the *E. coli* expression strain BL21-CodonPlus(DE3)-RIL (Stratagene). Purified PARG was tested for enzymatic activity on auto-PARylated PARP1. For auto-PARylation, 2 µg PARP1 (Enzo Life Sciences) was incubated in 10 µl of 50 mM Tris-HCl pH 8.0, 100 mM NaCl and 10 µM $NAD^+$ for 30 min at 37°C. To test for PAR glycohydrolase activity, 2 µg PARG was incubated with 0.5 µg auto-PARylated PARP1 in 20 µl 1x PARG buffer (50 mM Tris pH 8.0, 2 mM $MgCl_2$, 1 mM DTT, 100 mM NaCl ) at 30°C overnight. The reaction products were analyzed for PARylation by dot blot (see below).

*PARP1*

Human full-length *PARP1* cDNA (pCMV3-HA-PARP1, SinoBiological) was inserted into a pFastBac-vector, encoding an N-terminal $His_6$-MBP-HRV-3C-tagged fusion protein. $His_6$-MBP-HRV-3C-PARP1 was produced in 1.2 l SF9 insect cells using the Baculovirus expression system. PARP1 was purified essentially as described[37]. The His-MBP-tag was removed using 3C protease over night at 4°C prior to the Heparin chromatography step, and the final purified protein was stored in 25 mM HEPES-KOH pH 7.4, 300 mM NaCl, 1 mM EDTA, 1 mM DTT, 10% glycerol.

**Nuclease and PARG treatments**

PARG treatment of genomic DNA was done with 0.5 µg purified PARG per 1 µg of DNA in 1x PARG buffer (50 mM Tris-HCl pH 8.0, 2 mM $MgCl_2$, 1 mM DTT, 100 mM NaCl) at 30°C overnight, followed by Proteinase K (Qiagen, 10 µg per 1µg DNA, 4 h, 50°C) digestion and phenol/chloroform extraction of DNA. DNase I digestion was performed with 1 U DNase I (ThermoFisher) per 1 µg of DNA or PAR polymer (R&D Systems) for 1h at 37°C according to manufacturer's instruction, and followed by phenol/chloroform extraction. For triple-RNase treatment, genomic DNA/PAR polymer was consecutively incubated with 10 µg RNase A (ThermoFisher) per 1 µg of DNA/PAR polymer in TE buffer supplemented with 300 mM NaCl, 4 U ShortCut RNase III (NEB) per 1 µg of DNA/PAR polymer, and 2.5 U RNase H (NEB) per 1 µg DNA/PAR polymer according to supplier's instructions for 1h at 37°C. DNA/PAR polymer was phenol/chloroform extracted after each single reaction. Nuclease S1 treatment was done with 1 U Nuclease S1, respectively, per 1 µg genomic DNA for 1h at 37°C, and with phenol/chloroform extraction after the reaction. For Southwestern analysis, DNA was treated with 20 U EcoRI (NEB) or 20 U MseI (NEB) per 1 µg of DNA according to manufacturer's instructions for 1h at 37°C and subsequent phenol/chloroform extraction.

**Dot blot**

Genomic DNA was isolated using the Blood & Cell Culture Kit (Qiagen) according to manufacturer's instruction, and with overnight Proteinase K treatment, dissolving of DNA in presence of 20 µg/ml RNase A and a final phenol/chloroform extraction. Genomic DNA from human adult normal tissues was purchased from Amsbio (brain, HG-201; thymus, HG-702; heart, HG-801; skeletal muscle, HG-102; liver, HG-314; pancreas, HG-313; kidney, HG-901; spleen, HG-701; testis, HG-401; placenta, HG-413). Human tissue DNA was treated overnight with Proteinase K (0.3 mg/ml final concentration) followed by incubation with RNaseA (20 µg/ml final concentration) and phenol/chloroform extraction. Plasmid DNA (pEGFP-C1, Clontech) was prepared by QIAprep Spin Miniprep Kit. DNA was denatured for 10 min at 95°C in 500 µl 2x SSC and loaded on a 6x SSC equilibrated Hybond®-N+ Blotting Membrane using a Bio-Dot microfiltration unit (Bio-Rad) according to manufacturer's instructions. DNA was crosslinked to the membrane with a Stratalinker UV Crosslinker using the auto crosslink mode. DNA loading on membrane was visualized with 0.1% methylene blue in 0.5 M sodium acetate. For immunodetection of DNA PARylation, membranes were washed twice in TBST, blocked with 5% non-fat dry milk, incubated overnight with a 1:5000 dilution of an anti-poly (ADP-ribose) mouse monoclonal antibody (Trevigen, 4335-MC-100) or anti-pan-ADP-ribose biding reagent

(Merck, MABE1016), washed 3x with TBST, incubated for 1h at room temperature with a 1:5000 dilution of a goat anti-mouse IgG-HRP conjugate (Dianova, 115-035-146) or a goat anti-rabbit IgG-HRP conjugate (Dianova, 111-035-144) and washed 3x with TBST. Chemiluminescence was induced with SuperSignal West Femto ECL solution (ThermoFisher) according to manufacturer's instructions and recorded on a Bio-Rad ChemiDoc imaging system.

**dsDNA immunoprecipitation**

Genomic DNA of mESC was prepared with Blood & Cell Culture Kit (Qiagen) as described above. DNA immunoprecipitation was performed as described below using mouse anti-ds DNA antibody [35I9 DNA] (Abcam ab27156) and mouse IgG control (I8765). After final resuspension, 5 µl of each sample was subjected to dot blot analysis as described above.

**Southwestern blotting assay**

Genomic DNA was prepared with Blood & Cell Culture Kit (Qiagen) as described above and separated by electrophoresis on a 1% agarose gel (1 µg DNA per sample) followed by capillary blotting to a Hybond-N+ Membrane (GE Healthcare) using 20x SSC as blotting buffer. Crosslinking of DNA and immune-detection of DNA PARylation was as described above.

**Quantification of ribosyl-adenosine (R-Ado) by LC-MS/MS**

*Generation of $^{15}$N-labeled ribosyl-adenosine*

Stable isotope labeling of PAR polymers was performed by two consecutive enzymatic reactions essentially as described[19]. Briefly, $^{15}N_5$-$NAD^+$ was generated in a reaction containing 2mM $^{15}N_5$-labeled ATP (Silantes), 2 mM β-nicotinamide mononucleotide (N3501, Sigma) and 0.1 µg/µl nicotinamide-nucleotide adenylyltransferase (ProSpec, ENZ-1002) in 25 mM Tris-HCl, pH 7.5, 20 mM $MgCl_2$ for 1h at 37 °C. Isotopically labeled $NAD^+$ was used for PARP1 auto-PARylation as described above followed by phenol/chloroform extraction of the labeled PAR polymers. Purified $^{15}$N-PAR chains were degraded with nuclease P1 (NP1, Roche), snake venom phosphodiesterase (SVP, Worthington) and fast alkaline phosphatase (FastAP, Fermentas). The resulting $^{15}N_5$-ribosyl-adenosine ($^{15}N_5$-R-Ado) and $^{15}N_5$-diribosyl-adenosine ($^{15}N_5$-2R-Ado) were separated on an Agilent 1290 Infinity Binary LC system (Agilent Technologies) using a ReproSil 100 C18 column (Jasco). Isotopically labeled R-Ado and 2R-Ado were identified by analytical HPLC in tandem with triple quadruple mass spectrometry (Agilent 6490, Agilent Technologies) and purified by preparative HPLC. Concentrations of labeled R-Ado and 2R-Ado were experimentally determined by LC-MS/MS with defined concentrations of unlabeled reference R-Ado and 2R-Ado obtained from degraded PAR polymers (R&D systems, molar extinction coefficient 13.5 $mM^{-1}cm^{-1}$, assumed average length of 150 monomers).

*Genomic DNA preparation and LC-MS/MS analysis*

Genomic DNA was prepared with Blood & Cell Culture Kit (Qiagen) as described above but without a final phenol/chloroform extraction. PARG- and DNaseI-treated DNA was subjected to a second column purification. Mouse organ tissues were obtained from 7-8 weeks old female C57BL/6J mice. Tissues were homogenized with an Ultra-TURRAX disperser (IKA) prior to DNA preparation. About 10 µg of DNA was degraded to nucleosides with NP1 (Roche), SVP (Worthington) and FastAP (Fermentas). An equal volume of isotopic standard mixture $^{15}N_5$-dG (Silantes), $^{15}N_5{}^{13}C_{10}$-dA (Silantes) and self-synthesized $^{15}N_5$-R-Ado and $^{15}N_5$-2R-Ado (see above) was added to the DNA samples and ~10 µg of total DNA was injected for LC-MS/MS analysis. Quantitative analysis was performed on an Agilent 1290 Infinity Binary LC system (Agilent Technologies) using ZORBAX SB-C18 column (Agilent Technologies, 5 mm, 2.1×50 mm) coupled to an Agilent 6490 triple quadrupole mass spectrometer. Quantification of R-Ado and R2-Ado by LC-MS/MS was performed according to the published protocol[19] with specific changes: Elution was performed with 5 mM ammonium acetate pH 6.9 and acetonitrile (ACN), the flow was first linearly increased from 0.3 ml/min to 0.38 ml/min in 0 - 10.5 min, then switched to 0.5 ml/min for 10.5 - 14.5 min, and 0.3 ml/min for 14.5 - 15.5 min. The column was kept at 30 °C. The gradient was: 0 - 3 min, 0% ACN; 3 - 7.5 min, 0 - 5% ACN; 7.5 - 10.5 min, 5 % ACN, 10.5 -12.5 min 5 - 50% ACN; 12.5 - 15.5 min, 0% ACN. The MS source-dependent parameters were as follow: gas temperature 110 °C, gas flow 19 l/min ($N_2$), Nebulizer 25 psi, sheath gas heater 375 °C, sheath gas flow 11 l/min ($N_2$), capillary voltage 2000 V (positive mode), nozzle voltage 0 V, fragmentor voltage 300 V, high pressure RF 150 V and low pressure RF 60 V. Compound dependent parameters are listed in Extended Data Table 2. Note, in none of the gDNA samples we observed signals for 2R-Ado, the expected product for branched PARylation. R-Ado quantification is shown over dN as calculated from total dG and dA signals. To calculate limit of detection (LOD) of R-Ado molecules per genome, the LOD of the method ($3 \times 10^{-17}$ moles or $1.8 \times 10^{7}$ molecules of R-Ado) was divided by the amount of genomes in 10 µg injected DNA, i.e. ~$3.2 \times 10^{6}$ mouse or human genomes (3.1 pg average weight per genome).

**LC-MS/MS for base PARylation**

*Generation of 83mer ssDNA substrates with isotopically labeled nucleotides*

Asymmetric PCR[38] was used to generate 83mer ssDNA substrates with heavy isotope labeled nucleotides. Each PCR reaction contained 1000 nM forward primer, 50 nM reverse primer, a 83mer synthetic template DNA (primer and template sequences are listed in Extended Data

Table 1) and a dNTP-mixture in which one unlabeled dNTP was substituted with the respective heavy isotope-labeled dNTP ($^{15}N_5$-dGTP, $^{15}N_3$-dCTP, $^{15}N_2$-TTP, $^{13}C_{10}$-dATP or $^{15}N_5$-dATP, Silantes). Amplification was performed in 50 cycles. Subsequently, excess primers and dNTPs were removed by Amicon Ultra-0.5 ml centrifugal filter units (Millipore) according to manufacturer's instructions. The concentrated ssDNA was used for *in vitro* ADP-ribosylation and LC-MS/MS analysis as described below. Note, the 83mer ssDNA substrates contain a mixture of the respective heavy and natural nucleotide due to usage of unlabeled primer sequences.

*DNA preparation and LC-MS/MS analysis*

Degradation of *in vitro* PARylated DNA and LC-MS/MS conditions were essentially as described above for R-Ado quantification except that the MS source-dependent parameters for capillary voltage were 2200 V (positive mode), and high pressure RF was 130 V. When indicated, DNA was not heat-denatured at 95°C for 5 min prior to degradation, and 100 nM pentostatin (Sigma) was added to the degradation mixture. Ribosyl-dA or ribosyl-dI quantification is shown over total dA. Compound dependent parameters are listed in Extended Data Table 2.

*Enrichment of ribosyl-deoxyinosine (R-dI) from gDNA*

Male adult pig liver and kidneys dissected from 8-12 weeks old male C57BL/6J mice were homogenized with a Dounce homogenizer or an Ultra-TURRAX disperser (IKA). Genomic DNA was prepared with Blood & Cell Culture Kit (Qiagen) as described above omitting the final phenol/chloroform extraction but including an additional RNase A treatment (50 µg RNase A per 300 µg gDNA at 0.3 mg/ml) in 2 mM Tris-HCl pH 7.5, 18 mM ammonium acetate for 30 min at 37°C followed by ethanol precipitation. Degradation of ssDNA was performed with 1400 units of Nuclease S1 per 1 mg of gDNA in 5 mM ammonium acetate pH 5.6, 0.2 mM $ZnCl_2$ for 60 min at 37 °C using a total amount of 5 and 2.5 mg of pig liver and mouse kidney DNA, respectively, at 0.7 mg/ml. Degraded ssDNA was separated from undigested dsDNA by an Amicon ®Ultra-0.5 ml 30 kDa filter unit (Millipore). The ssDNA collected from the flow through was hydrolyzed with 0.4 U of NP1, 2 U of SVP and 20 U of FastAP per 1 mg of starting gDNA amounts followed by enzyme removal through an Amicon ®Ultra-0.5ml 10 kDa filter unit (Millipore). The flow through was concentrated ~20x in a Concentrator plus (Eppendorf) at 4 °C. R-dI was enriched on an Agilent 1290 Infinity Binary LC system (Agilent Technologies) using a 250 mm × 4,6 mm ReproSil 100 C18 3 µm (Jasco) by sequential runs with 0.8 mg of starting DNA amount per injection. LC was performed with 5 mM ammonium acetate pH 6.9 and acetonitrile (ACN), the flow was at 0.5 ml/min for 0 - 40 min, then gradually increased to 1 ml/min for 40-50 min, gradually decreased to 0.5 ml/min for 50-55 min. The gradient was: 0 -

15 min, 0% ACN; 15 - 35 min, 0 - 15% ACN; 35 - 40 min, 15 - 50 % ACN, 40 - 45 min 50% ACN; 45 - 55 min, 0% CAN at 30 °C. The collection window was from 29.9-30.9 min as determined empirically from *in vitro* generated R-dI, which peaks at ~30.5 min. The collected fractions were pooled, concentrated to ~20 µL and analyzed in a single analytical LC-MS/MS run using the same column and LC conditions as for R-dI purification. Compound dependent parameters are listed in Extended Data Table 2.

**Quantitative real time PCR (qPCR)**

Quantitative real time PCR was performed on a LightCycler 480 (Roche) in technical duplicates using the Universal ProbeLibrary technology (Roche) in combination with the supplier's LightCycler 480 Probes Master. Quantitative analysis was performed with LightCycler 480 software (Roche). For reverse transcription-coupled (RT) qPCR, RNA was isolated using the RNeasy mini kit (Qiagen) following the manufacturer's instructions. cDNA synthesis was performed with SuperScript II Reverse Transcriptase (ThermoFisher). Primer sequences and hydrolysis probe numbers are listed in Extended Data Table 1.

**H2A.X chromatin immunoprecipitation (ChIP)**

DIvA cells were treated with 300 nM 4-OHT (Sigma, H6278) or mock treated for 24 h before harvesting. ChIP assay was carried out as described[24] using 200 µg of sonicated chromatin and 2 µg mouse monoclonal anti-phospho-Histone H2A.X antibody (Millipore, 05-636-I) per sample. ChIP efficiencies were calculated using the percent input method after qPCR and are shown as fold change over mock treatment.

**PARylated DNA immunoprecipitation (PAR-DIP)**

Genomic DNA was prepared with Blood & Cell Culture Kit (Qiagen) as described above. DNA was sonicated in a Bioruptor (Diagenode) according to manufacturer's instructions to generate fractions between 300 and 500 bp followed by phenol/chloroform extraction and ethanol precipitation. DNA immunoprecipitation was performed overnight at 4 °C with 5 µg of sonicated DNA and 5 µg of anti-poly (ADP-ribose) mouse monoclonal antibody (Trevigen, 4335-MC-100) in 500 µl of 1X IP buffer (10 mM Na-phosphate pH 7.0, 140 mM NaCl, 0.05% Triton X-100). As input 1-10% of sonicated DNA was kept separately. Antibodies were captured by 2h incubation with Dynabeads Protein G (ThermoFisher, 40 µl per IP sample) pre-washed with 0.1 % BSA. Following a three-time wash of beads with 700 µl IP buffer, Proteinase K digestion (0.3 mg/ml final concentration) was performed on beads and input samples for 3h at 50 ºC. Finally, DNA was phenol/chloroform extracted, ethanol precipitated and resuspended in 20 µl nuclease-free $H_2O$. Target regions of γH2AX-associated and -not associated AsiSI sites (Extended Data Table 1) were chosen based on a previous analysis[24].

PAR-DIP efficiencies were calculated using the percent input method after qPCR and are shown as fold change over mock treatment.

**Immunofluorescence**

Diva cells grown on coverslips and treated or mock treated for 24h with 300 nM 4-OHT were fixed in 4% formaldehyde solution, neutralized with 200 mM Glycine, permeabilized in 0.2% Triton X-100 and blocked with 5% BSA. Primary antibody incubation was done overnight at 4 °C using a mouse monoclonal anti-phospho-Histone H2A.X antibody (Millipore, 05-636-I, 1:300 dilution in 1% BSA) followed by secondary antibody incubation for 2h at room temperature using a goat anti-mouse IgG Alexa Fluor 488 conjugate (ThermoFisher, A-11029, 1:500 dilution in 1% BSA). DNA was stained 10 min in 0.5 µg/ml DAPI and coverslips mounted with ProLong Gold (ThermoFisher). Images were acquired using a Leica TCS SP5 confocal microscope.

***In vitro* ADP-ribosylation assay**

End-labeling of DNA was performed with 200 nM oligonucleotide with [γ-32P]ATP (PerkinElmer) and T4 polynucleotide kinase (NEB) according to the manufacturer's instructions. 100 nM labeled DNA was hybridized in SSC buffer (150 mM NaCl and 15 mM trisodium citrate) with 100 nM complementary DNA or RNA oligonucleotides (Extended Data Table 1). Unincorporated [γ-32P]-ATP was removed by G-25 Quick Spin columns (GE Healthcare). ADP-ribosylation assay was performed with 1 nM single-stranded or duplex oligonucleotides, 1 mM $NAD^+$, 20 nM recombinant PARP1 and 200 nM olaparib if indicated in ADPR buffer (20 mM HEPES-KOH, pH 7.6, 50 mM KCl, 1 mM DTT, 100 µg/ml BSA) for 60 min at 37 °C followed by treatment with 5 U alkaline phosphatase (CIP, NEB) if indicated. To test for PAR chain addition, ADP-ribosylation assay was performed in presence of 0.1 – 10 µM of PAR polymer (Trevigen, molar extinction coefficient 13.5 $mM^{-1}cm^{-1}$). The reaction was stopped by addition of 50 ng/µl Proteinase K for 30 min at 56 °C. Reaction products were denatured in 1x Novex TBE-Urea Sample Buffer (ThermoFisher) and analyzed on 10% Novex TBE-Urea Gels (ThermoFisher) according to manufacturer's instruction. Phosphorimaging was performed on a Typhoon FLA 9500 (GE Healthcare).

For LC-MS/MS analysis, ADP-ribosylation was performed with 200 nM DNA substrate, 2 mM $NAD^+$, 2 µM recombinant PARP1, and 20 µM olaparib or 100 nM pentostatin (Sigma) if indicated in ADPR buffer for 60 min at 37 °C followed by phenol/chloroform extraction and ethanol precipitation. PARG treatment was done with 200 nM *in vitro* PARylated DNA substrate and 0.2 ng/µl PARG in PARG buffer for 2 h at 30 °C. To remove free PAR chains PARG treatment was followed by DNA cleanup with DNA clean and concentrator 5 (Zymo Research) according to manufacturer's instructions. Control reactions were done with PARP1 or PARG heat-denatured at 70 °C for 10 min.

**Statistics**

Data presented as bar diagrams are displayed as arithmetic mean, error bars represent standard deviation (s.d.) of the indicated replicates with propagation of error if required after data normalization. Statistical analysis was carried out with Student's t-test as indicated in the figure legends. For multiple comparisons adjusted p-values were calculated by Dunnett's test using GraphPad Prism. Significances are displayed as *$p<0.05$, **$p<0.01$, ***$p<0.005$. n.s., not significant.

**Data and code availability**

All data are available from the corresponding author upon request.

**Methods references**

35. Dawlaty, M. M. *et al.* Loss of Tet enzymes compromises proper differentiation of embryonic stem cells. *Developmental cell* **29,** 102–111; 10.1016/j.devcel.2014.03.003 (2014).

36. Amé, J.-C., Héberlé, É., Camuzeaux, B., Dantzer, F. & Schreiber, V. Purification of Recombinant Human PARG and Activity Assays. *Methods in molecular biology (Clifton, N.J.)* **1608,** 395–413; 10.1007/978-1-4939-6993-7_25 (2017).

37. Langelier, M.-F., Planck, J. L., Servent, K. M. & Pascal, J. M. Purification of human PARP-1 and PARP-1 domains from Escherichia coli for structural and biochemical analysis. *Methods in molecular biology (Clifton, N.J.)* **780,** 209–226; 10.1007/978-1-61779-270-0_13 (2011).

38. Yunusov, D. *et al.* Kinetic capillary electrophoresis-based affinity screening of aptamer clones. *Analytica chimica acta* **631,** 102–107; 10.1016/j.aca.2008.10.027 (2009).

### Extended Data Figures

### Extended Data Fig. 1

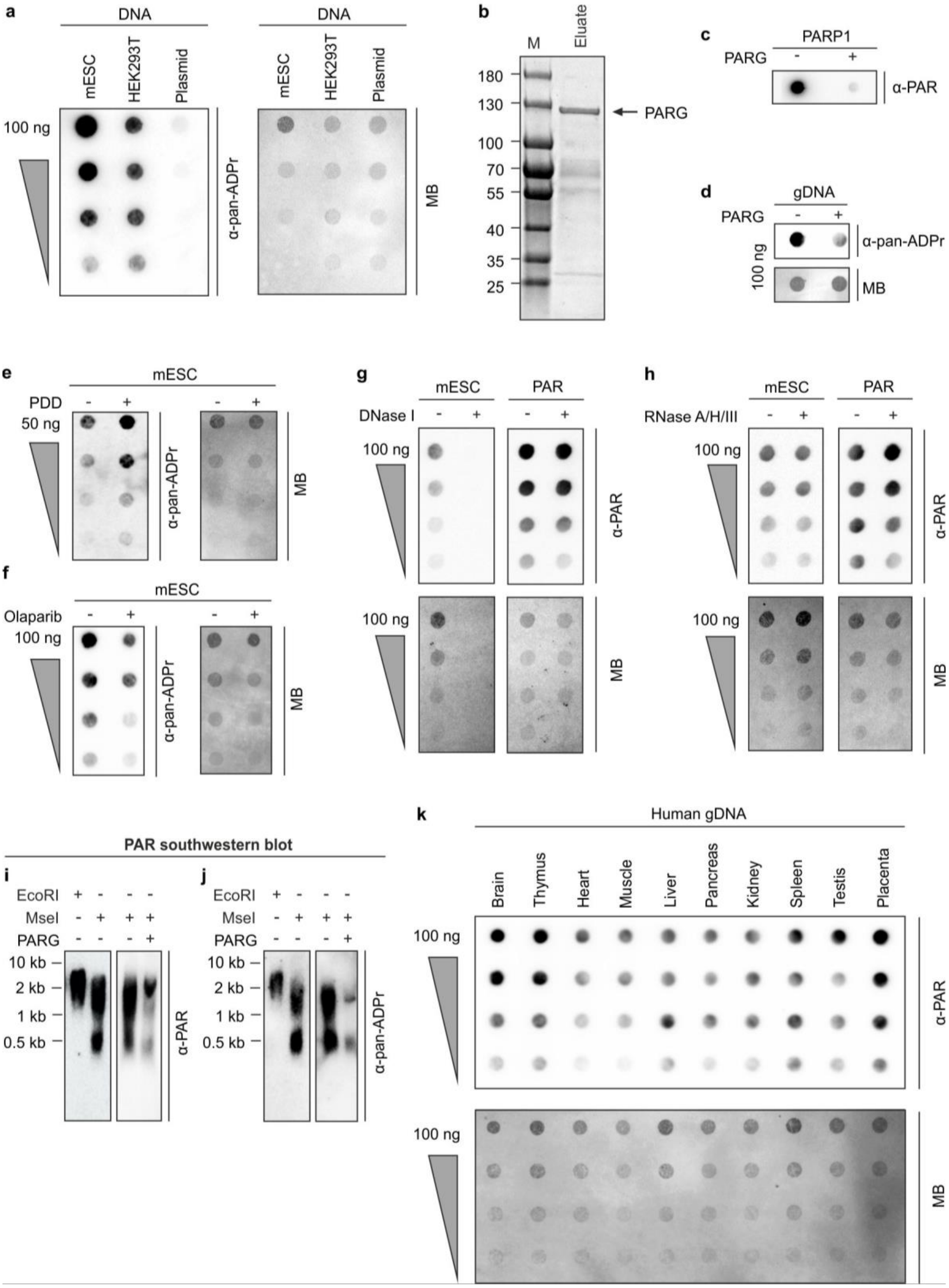

**Extended Data Fig. 1: DNA PARylation in mouse and human tissues.**

**a,** Dot blot analysis for PARylation of mESC and HEK293T serially (2x) diluted genomic DNA as in Fig. 1a but using anti-pan-ADP-ribose binding reagent. Plasmid DNA served as negative control. MB, methylene blue staining.

**b**, SDS-PAGE analysis of the purified full-length PAR glycohydrolase (PARG[36], arrow) used in the study. Calculated molecular weight for PARG is 112 kDa. Molecular weight of marker proteins (M) is indicated on the left in kDa. Eluate, supernatant post treatment with PreScission protease.

**c-h**, Dot blot analysis for PARylation employing either anti-poly (ADP-ribose) mouse monoclonal antibody (α-PAR) or anti-pan-ADP-ribose binding reagent (α-pan-ADPr) of (**c**) autoPARylated PARP1 treated with PARG as indicated, (**d**) mESC DNA treated with PARG, (**e**) DNA purified from mESCs treated with the PARG inhibitor PDD00017273, (**f**) DNA purified from mESCs treated with the PARP inhibitor olaparib, (**g**) genomic mESC DNA (left) and PAR polymer (right) treated with DNase I, (**h**) genomic mESC DNA (left) and PAR polymer (right) treated with RNase A, RNase H and RNase III, MB, methylene blue.

**i-j**, Southwestern blot analysis for PARylation of HEK293T DNA treated with EcoRI, MseI, and PARG as indicated and utilizing anti-poly (ADP-ribose) mouse monoclonal antibody (α-PAR, **i**) or anti-pan-ADP-ribose binding reagent (α-pan-ADPr, **j**). Length of marker DNA is shown on the left.

**k**, Dot blot analysis for PARylation of DNA from the indicated adult human tissues. gDNA was serially diluted (2x). MB, methylene blue.

**Extended Data Fig. 2**

a

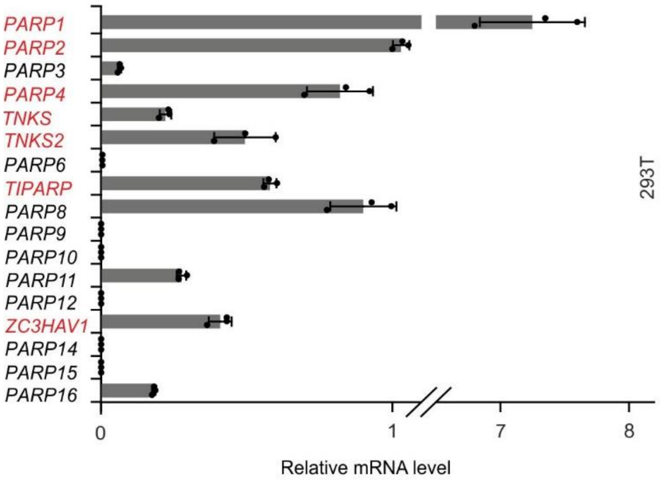


b

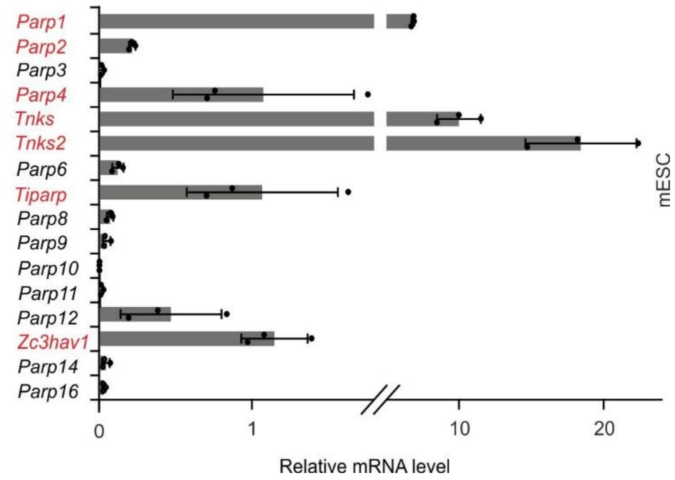


**Extended Data Fig. 2: PARP candidates for DNA PARylation selected by their expression levels.**

**a**, Expression analysis of *PARP* genes in 293T cells (s.d., n = 3 biological replicates). *PARP genes* commonly high expressed in 293T and mESCs are highlighted in red.

**b**, Expression analysis of *Parp* genes in mESC (s.d., n = 3 biological replicates). *Parp genes* commonly high expressed in 293T and mESCs are highlighted in red.

**Extended Data Fig. 3**

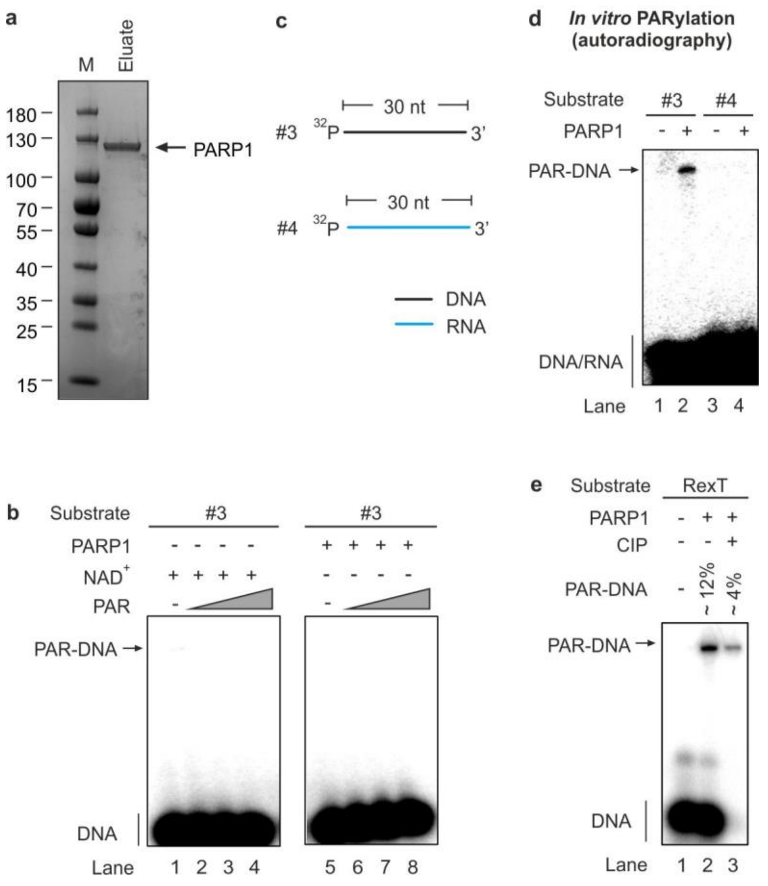


**Extended Data Fig. 3: PARP1 preferentially PARylates ssDNA *in vitro*.**

**a,** SDS-PAGE analysis of purified full-length PARP1 (arrow) used in the study. Calculated molecular weight for PARP1 is 113 kDa. Molecular weight of marker proteins (M) are indicated on the left in kDa. Eluate, PARP1 pooled fraction from gel filtration chromatography.

**b**, Free PAR chains are not conjugated to DNA. Substrate #3 was incubated with increasing amounts of free PAR chains in presence of $NAD^+$ or PARP1 as indicated followed by denaturing PAGE and autoradiography of the reaction products. Note the absence of PARylated DNA products, indicating that free PAR chains are not attached to substrate ssDNA either spontaneously or by PARP1.

**c**, Scheme of ssDNA and ssRNA substrates used for *in vitro* PARylation assays. The 30 nt DNA strand is designated as standard oligo in the main text. $^{32}P$, 5'-phosphate with $^{32}P$-label.

**d-e**, Denaturing PAGE and autoradiography of reaction products from PARylation assays in presence of PARP1, with post reaction CIP-treatment and substrates as indicated. (**d**), PARP1 does not PARylate ssRNA. (**e**), PARP1 PARylates ssDNA RexT both internally (8%) and terminally (4%).

### Extended Data Fig. 4

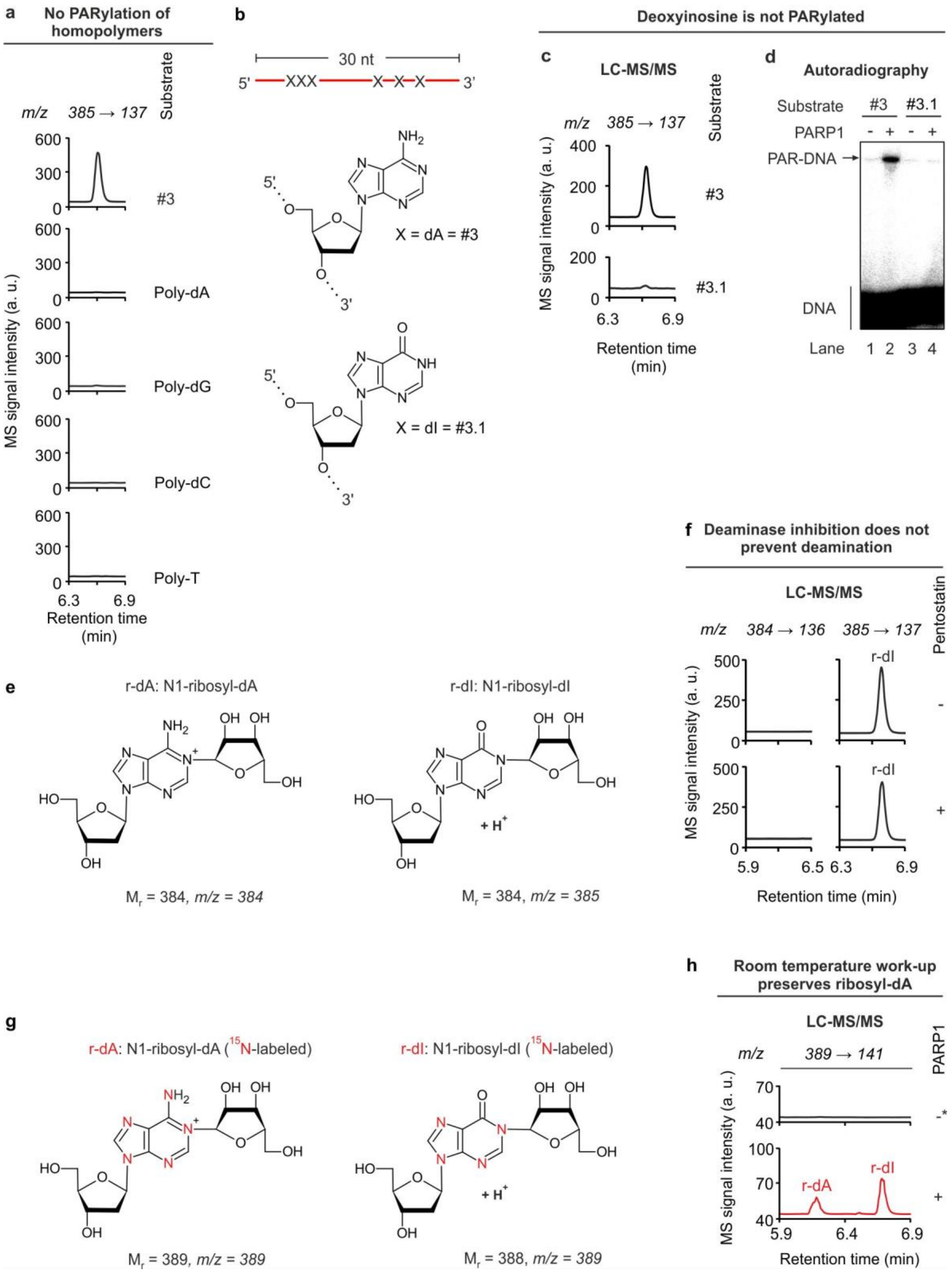


**Extended Data Fig. 4: Identification of the PARylated base by LC-MS/MS analyses.**

**a**, LC-MS/MS chromatograms of reaction products from *in vitro* PARylation assays with the indicated substrates and human PARP1. Mass transition corresponds to shift expected for loss of a deoxyribose + ribose (*m/z* 385 → 137) from the parental molecule 'nucleoside 385'.

**b**, Scheme of the 30mer ssDNA standard oligo used for *in vitro* PARylation assays. In substrate #3 'X' represents dA, in substrate #3.1 'X' is dI. Structures of dA and dI are shown below.

**c**, LC-MS/MS chromatogram for 'nucleoside 385' in reaction products from PARylation assay with the substrates indicated.

**d**, Autoradiography of denaturing PAGE of reaction products from PARylation assay with the indicated substrates.

**e**, Structure, relative molecular weight and *m/z* ratio for N1-ribosyl-dA (left) and N1-ribosyl-dI (right).

**f**, LC-MS/MS chromatograms of reaction products from *in vitro* PARylation assay in presence of substrate #3 and human PARP1. PAR reaction and enzymatic degradation of reaction products for mass spec analysis was done in presence or absence of the deaminase inhibitor Pentostatin as indicated. Products were scanned for signals with *m/z* transitions expected for N1-ribosyl-dA (r-dA, 384 → 136, left) or N1-ribosyl-dI (r-dI, 385 → 137, right)

**g**, Structure, relative molecular weight and *m/z* ratio for $^{15}N$-labeled N1-ribosyl-dA (left) and $^{15}N$-labeled N1-ribosyl-dI (right). Heavy isotope labeled nitrogens are highlighted in red.

**h**, LC-MS/MS chromatograms of reaction products from *in vitro* PARylation assay with the $^{15}N_5$-dA-labeled 83mer ssDNA oligo in presence of native and denatured (-*) PARP1 as indicated. Unlike for standard protocol, sample denaturation at 95 °C was omitted before mass spec analysis to avoid deamination. Samples were screened for signals with the same *m/z* transition of 389 → 141 as expected for both $^{15}N_5$-labeled N1-ribosyl-dA and $^{15}N_4$-labeled N1-ribosyl-dI, but distinguished by different retention times.

### Extended Data Fig 5

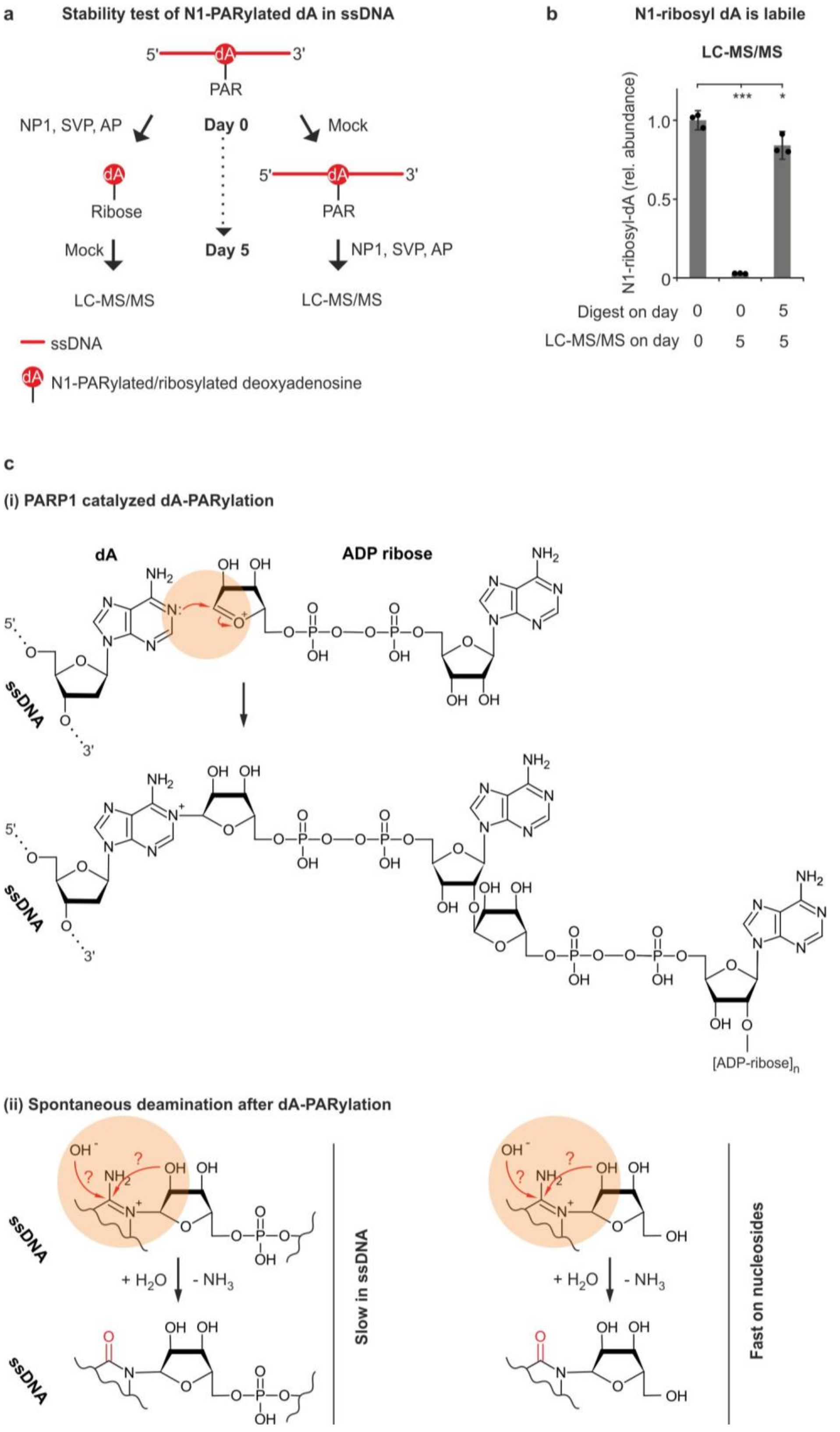


**Extended Data Fig. 5: Stability of PARylated-dA in ssDNA and proposed reaction mechanisms.**

**a**, Outline of stability test for PARylated dA within ssDNA. An *in vitro* PARylated 83mer oligonucleotide is either degraded to single nucleosides by combined treatment with nuclease P1, snake venom phosphodiesterase and alkaline phosphatase (NP1, SVP, AP) or mock-treated on day 0. Samples are stored at 4 °C for 5 days followed by mock treatment of single nucleosides and degradation of the 83mer with NP1, SVP, AP and subsequent LC-MS/MS analysis.

**b**, LC-MS/MS quantification of N1-ribosyl-dA in samples as described in a. As reference, the PARylated 83mer was degraded and analyzed on day 0, and arbitrarily set to 1. s.d., n = 3 biological replicates; *$P < 0.05$ and ***$P < 0.005$ by Dunnett's test. Note, while we readily detect a signal for N1-ribosyl-dA on samples that were kept non-degraded for 5 days, the signal was erased in samples that were degraded immediately and stored for 5 days at -20°C. Moreover, samples that were degraded and analyzed directly after *in vitro* PARylation show marginally higher (~20%) N1-ribosyl-dA signals compared to samples stored for 5 days and digested only before LC-MS/MS. Hence, deamination is slow on PARylated-dA in intact DNA but vastly accelerates on ribosylated dA-nucleosides.

**c**, Proposed reaction mechanisms. (i), PARP1 catalyzes a nucleophilic attack of N1 from dA to C1 of the oxonium ion of ADP ribose, followed by PARylation. (ii), Spontaneous deamination is slow on PARylated dA within ssDNA but fast on ribosyl-dA nucleosides, and presumably proceeds via hydrolytic attack at C6 by free hydroxyl ions or hydroxyl groups of the attached ribose as indicated by two alternative arrows.

**Extended Data Tables**

**Extended Data Table 1: Oligonucleotides used in this study.**

PAR-DIP- and ChIP-qPCR primers (IDT) and UPL probe numbers (Roche)

| **Target** | **Direction** | **Sequence (5’→3‘)** | **Probe #** |
|---|---|---|---|
| human Chr1: 88992918 | Forward<br>Reverse | CACCCCTCATCCAGTCTCTT<br>ATTGACTCTGGGGTTTGCAC | 162 |
| human Chr6: 89638472 | Forward<br>Reverse | CAGCTCAGACCCAGGTAAGC<br>GGTGGACATGGCTGCTTC | 13 |
| human Chr22: 20496025 | Forward<br>Reverse | CCTCGGTCCTACCCTCATTC<br>CGGGGAGGGAGATAGTGAG | 51 |
| human Chr1: 3186484 | Forward<br>Reverse | CCTGGGATGGAGGACTCTC<br>GCCCCTGTCTACATTCCTGA | 3 |

RT-qPCR primers (IDT) and UPL probe numbers (Roche)

| **Gene** | **Direction** | **Sequence (5’→3‘)** | **Probe #** |
|---|---|---|---|
| mouse *Gapdh* | Forward<br>Reverse | AGCTTGTCATCAACGGGAAG<br>TTTGATGTTAGTGGGGTCTCG | 9 |
| mouse *Parp1* | Forward<br>Reverse | AGGCCGCCTACTCTATCCTC<br>GATTCACTGCTGCCTTGAGA | 40 |
| mouse *Parp2* | Forward<br>Reverse | TCGTCCTTCAAGAGCGATG<br>CCAAGAATTACTGCACTATGAAGATG | 97 |
| mouse *Parp3* | Forward<br>Reverse | TCAGTGCTGTGCAATAGACTCTTA<br>GGGGCTTAGGAAGATGTGTG | 2 |
| mouse *Parp4* | Forward<br>Reverse | TGCAACAGCGCAGTCTCTTA<br>TGCTCCACAGCTTTCAGTTG | 34 |
| mouse *Tnks (Parp5a)* | Forward<br>Reverse | CCCACACACAAAGACAGATCA<br>GCCATTTTCATGGTGCTGA | 4 |
| mouse *Tnk2 (Parp5b)* | Forward<br>Reverse | TCCAATTCACAAAGACAGATCG<br>TCCCAAGGTTACTCGACAAAA | 27 |
| mouse *Parp6* | Forward<br>Reverse | GGCTACGGTTTTCTCTCTCTCA<br>CCTTATTCGATGGCTGGAAA | 21 |
| mouse *Tiparp (Parp7)* | Forward<br>Reverse | CAGAACAGGGGGTTCCAAT<br>CCACTGTCCCACTGATGGTT | 31 |
| mouse *Parp8* | Forward | CGGTATTTTAACACCATCATCG | 75 |

| | | | |
|---|---|---|---|
| | Reverse | GGAATGCTTTTTGCACCATT | |
| mouse *Parp9* | Forward<br>Reverse | TCAGCGGCTGGACTCATAC<br>GAGAGTGCGGTCCACAGAC | 58 |
| mouse *Parp10* | Forward<br>Reverse | GAACCCTTCTAAGGGCAGGT<br>GGATCCTAATTGATCCCGAGT | 1 |
| mouse *Parp11* | Forward<br>Reverse | GGCGTTGGGAGTTCAGAGT<br>CAAACAAACAAACAAACAAAAACA | 2 |
| mouse *Parp12* | Forward<br>Reverse | AAAGGGCTTACTTGGCATTCT<br>TGCCATACACAAGGTTTTTCTG | 1 |
| mouse *Zc3hav1 (Parp13)* | Forward<br>Reverse | AGGTATTCTGTTTCATCACCAAGA<br>CTTCGGGGAGGCTGATCT | 3 |
| mouse *Parp14* | Forward<br>Reverse | TGTGAAACAAGGTGATTTGGA<br>GGGACATCTTCTGTCTTTTCTGA | 3 |
| mouse *Parp16* | Forward<br>Reverse | GCCAAATAGAGGGGGAGATG<br>GCTTGGACCCTTCTCCATAA | 1 |
| human *TBP* | Forward<br>Reverse | GAACATCATGGATCAGAACAACA<br>ATAGGGATTCCGGGAGTCAT | 87 |
| human *PARP1* | Forward<br>Reverse | ACTTCCTCCAGGACGTCTCC<br>TGTGCGCTAAGAACAACTCC | 22 |
| human *PARP2* | Forward<br>Reverse | ACCAAGAAAGCCCCACTTG<br>AGCCCGAATACAATCCTCAA | 59 |
| human *PARP3* | Forward<br>Reverse | AACTGGGTAATCGGAAGCTG<br>ATGATGCGGAGCCCACTA | 49 |
| human *PARP4* | Forward<br>Reverse | GAAGTGAACCTGGGACTATTGG<br>TCACAGACATTAACCATGTCTCTTATT | 53 |
| human *TNKS (PARP5A)* | Forward<br>Reverse | GGCAAACGTAAATGCAAAGG<br>AACATCCTTCCTTCCAAAACCT | 7 |
| human *TNKS2 (PARP5B)* | Forward<br>Reverse | GGGTTGTTTAGCCAGAGTGAA<br>TCAGCTCCGTGTTGTAACAAAT | 2 |
| human *PARP6* | Forward<br>Reverse | GCCCCATCTCCAGTATTTCC<br>CTCATCCTTGGAGGGCATC | 56 |
| human *TIPARP (PARP7)* | Forward<br>Reverse | GGAAATTCTTCTGTAGGGACCA<br>AATCAATCGAATGACAGACTCG | 58 |
| human *PARP8* | Forward<br>Reverse | ATTTGGATTGGGACATCAGC<br>ATTGTGCAGGCAATTGGATT | 86 |
| human *PARP9* | Forward<br>Reverse | GAACAACCTGACCCTCCAGA<br>TCATGTGGGTTTACAGAATTAACAA | 81 |

| | | | |
|---|---|---|---|
| human *PARP10* | Forward<br>Reverse | CCTTCTACGACACCCTGGAC<br>CCGGTACAGCTCATACTGCTG | 89 |
| human *PARP11* | Forward<br>Reverse | CCCCCTTTTCTATCAGTGCTT<br>TGCAGAGGAATAAGCTGATATGG | 14 |
| human *PARP12* | Forward<br>Reverse | GTCCCACAGGGGACTTCTG<br>TTTTCCGGATATGGTACAAACA | 63 |
| human *ZC3HAV1 (PARP13)* | Forward<br>Reverse | AGACCAACATTTGTGCCTCA<br>CACTGACGAGGTCTTTGCTG | 84 |
| human *PARP14* | Forward<br>Reverse | GTGTTCTTCTACCCGGAGGA<br>TTCCTTGCCATACCAACTCA | 29 |
| human *PARP15* | Forward<br>Reverse | CACCCGAACTTGTTCTTCCTA<br>CCATTTGCCTTTTCTTTACCTG | 53 |
| human *PARP16* | Forward<br>Reverse | CAGCTGCTGCGAGTGAAG<br>CAGGAGAGCTGGCTCGAA | 79 |

siRNA sequences (siGENOME SMARTpool, mouse and human, Dharmacon)

| **Name** | **Target sequence** |
|---|---|
| siControl | 1: UAGCGACUAAACACAUCAA<br>2: UAAGGCUAUGAAGAGAUAC<br>3: AUGUAUUGGCCUGUAUUAG<br>4: AUGAACGUGAAUUGCUCAA |
| human si*PARP1* | 1: GAAAGUGUGUUCAACUAAU<br>2: GCAACAAACUGGAACAGAU<br>3: GAAGUCAUCGAUAUCUUUA<br>4: GAUAGAGCGUGAAGGCGAA |
| human si*PARP2* | 1: AAGGAUUGCUUCAAGGUAA<br>2: ACAGCUAGAUCUUCGGGUA<br>3: GCCAGAGACAGGAGUCGAA<br>4: ACAAUUGGGAAGAUCGAGA |
| human si*PARP4* | 1: CAACUGAACCACUAUUUAA<br>2: GAGCAGUUCUGAAGUGAAA<br>3: GUGCACACAUAUAAUCUUA<br>4: CCACAGACUUUGAGGAUGA |
| human si*TNKS (PARP5A)* | 1: GAACAGAGAUGGAAAUACA<br>2: CAGAGUAUCUUAUCACUUA<br>3: GGAAGUAGCUGAAUAUCUU |

| | |
|---|---|
| | 4: CAACAGAGUUCGAAUAGUU |
| human si*TNKS2 (PARP5B)* | 1: GGAAAGACGUAGUUGAAUA<br>2: UAGCAUAACUCAAUUCGUA<br>3: AGACAGAUCUUGUUACAUU<br>4: AAUGUAAAUUGCCGCGAUA |
| human si*TIPARP (PARP7)* | 1: GGCCGUGACAGGAUAAUAA<br>2: GAAAGAGGUUCGAUUUAUG<br>3: AAGGCAAGCUACUCUCAUA<br>4: GGCAGAGUGUAUUCAAUGA |
| human si*ZC3HAV1 (PARP13)* | 1: GCACAUGGAUUCAGUAUGG<br>2: GCAAGCACAUGCAGAAGAA<br>3: GAACAAAGAGGAAUUAGCA<br>4: CAAAUAUUCUCAUGAGGUU |
| mouse si*Parp1* | 1: GGAGGAAGGUGUCAACAAA<br>2: UAAAGAAGCUGACGGUGAA<br>3: CAAAGUAUCCCAAGAAGUU<br>4: GAAAUAUCCUACCUCAAGA |
| mouse si*Parp2* | 1: GGGAAAGGCUCAUGUGUAU<br>2: GAAGGCGAGUGCUAAAUGA<br>3: GCAAGAAGAUGCGCACGUG<br>4: GGACUAUACUAUGACCUUG |

DNA and RNA oligonucleotides used for *in vitro* PARylation (IDT and Sigma)

| **Name** | **Sequence (5'→3')** |
|---|---|
| 30mer DNA_up (#3) | TGTGCAAACGCCGGCTTCCGCACAGAGGCC |
| 30mer RNA_low (#4) | GGCCUCUGUGCGGAAGCCGGCGUUUGCACA |
| 30mer DNA_low | GGCCTCTGTGCGGAAGCCGGCGTTTGCACA |
| 40mer DNA_low | CACGCAAGGTGGCCTCTGTGCGGAAGCCGGCGTTTGCACA |
| 30mer DNA_up_2'd-inosine (#3.1) | TGTGC[dI][dI][dI]CGCCGGCTTCCGC[dI]C[dI]G[dI]GGCC |
| RexT | GGAATTCCCCGCGCCAAATTTCTCTAAGTCTCCGCGCCAC |
| 83mer DNA | CTCCTCTGACTGTAACCACGCCGGTACGTTACGATACGATTA<br>CGTAATACGATTTCGAACCGGCATAGGTAGTCCAGAAGCCT |
| Poly-dA | AAAAAAAAAAAAAAAAAAAA |
| Poly-dG | GGGGGGGGGGGGGGGGGGGG |
| Poly-dC | CCCCCCCCCCCCCCCCCCCC |

| | |
|---|---|
| Poly-T | TTTTTTTTTTTTTTTTTTTT |

dI, 2’d-inosine

DNA oligonucleotides (IDT) used for asymmetric PCR

| **Purpose** | **Direction** | **Sequence (5’→3‘)** |
|---|---|---|
| Primer | Forward | CTCCTCTGACTGTAACCACG |
| | Reverse | AGGCTTCTGGACTACCTATGC |
| Template | | CTCCTCTGACTGTAACCACGCCGGTACGTTACGATACGATTA<br>CGTAATACGATTTCGAACCGGCATAGGTAGTCCAGAAGCCT |

**Extended Data Table 2: MRM transitions used in this study.**

| Nucleoside | Precursor ion (*m/z*) | Fragment ion (*m/z*) | Collision energy | Cell accelerator voltage |
|---|---|---|---|---|
| dA | 252 | 136 | 6 | 8 |
| $^{15}N_5{}^{13}C_{10}$-dA | 267 | 146 | 6 | 8 |
| dG natural isotopologue (+1) | 269 | 153 | 6 | 5 |
| $^{15}N_5$-dG | 273 | 157 | 6 | 5 |
| R-Ado (1)(qualitative) | 400 | 136 | 6 | 8 |
| R-Ado (2)(quantitative) | 400 | 268 | 5 | 8 |
| $^{15}N_5$- R-Ado | 405 | 141 | 6 | 8 |
| 2R-Ado | 532 | 136 | 6 | 8 |
| $^{15}N_5$- 2R-Ado | 537 | 141 | 6 | 8 |
| N1-ribosyl-dI (‘nucleoside 385’) (1) | 385 | 137 | 17 | 5 |
| N1-ribosyl-dI (‘nucleoside 385’) (2) | 385 | 269 | 1 | 3 |
| N1-ribosyl-dA | 384 | 137 | 17 | 5 |
| $^{15}N_1$-‘nucleoside 385’ | 386 | 138 | 17 | 5 |
| $^{15}N_2$-‘nucleoside 385’ | 387 | 139 | 17 | 5 |
| $^{15}N_3$-‘nucleoside 385’ | 388 | 140 | 17 | 5 |
| $^{15}N_4$-ribosyl-dI, $^{15}N_5$-ribosyl-dA | 389 | 141 | 17 | 5 |
| $^{15}N_5$-‘nucleoside 385’ | 390 | 142 | 17 | 5 |
| $^{13}C_{10}$-ribosyl-dI | 395 | 142 | 17 | 5 |

MS 1 and 2 resolutions were set to unit for all ions. Note, all quantification were performed using stable-isotope dilution LC-MS/MS except for N1-ribosyl-dA or N1-ribosyl-dI due to lack of the respective isotopologue.
